# Unsupervised Multi-scale Segmentation of Cellular cryo-electron Tomograms with Stable Diffusion Foundation Model

**DOI:** 10.1101/2025.06.25.661425

**Authors:** Mostofa Rafid Uddin, Thanh-Huy Nguyen, H.M. Shadman Tabib, Kashish Gandhi, Min Xu

**Affiliations:** Carnegie Mellon University, Pittsburgh, PA 15213, USA; Bangladesh University of Engineering and Technology, Dhaka 1000, Bangladesh

**Author notes:** Corresponding Author: Min Xu. Equal Contributions (Thanh-Huy Nguyen, H.M. Shadman Tabib).

## Abstract

We introduce an unsupervised approach for segmenting multiscale subcellular objects in 3D volumetric cryo-electron tomography (cryo-ET) images, addressing key challenges such as large data volumes, low signal-to-noise ratios, and the heterogeneity of subcellular shapes and sizes. The method requires users to select a small number of slabs from a few representative tomograms in the dataset. It leverages features extracted from all layers of a Stable Diffusion foundation model, followed by a novel heuristic-based feature aggregation strategy. Segmentation masks are generated using adaptive thresholding, refined with CellPose to split composite regions, and then utilized as pseudo-ground truth for training deep learning models. We validated our pipeline on publicly available cryo-ET datasets of *S. Pombe* and *C. Elegans*cell sections, demonstrating performance that closely approximates expert human annotations. This fully automated, data-driven framework enables the mining of multi-scale subcellular patterns, paving the way for accelerated biological discoveries from large-scale cellular cryo-ET datasets.

## Introduction

Cells consist of multiple subcellular objects of diverse sizes, including large membranes and organelles, as well as smaller macromolecular complexes like ribosomes and proteasomes [1, 2, 3]. Thanks to cryo-electron tomography (cryo-ET), an *in situ* 3D imaging technology, it is now possible to visualize these objects of varying sizes together inside the cell in their native state [4]. In cryo-ET, a cellular specimen is rapidly frozen at cryogenic temperature without any purification or hampering of the specimen [5]. The vitreously frozen specimen is then placed under an electron microscope and gradually tilted around its tilt axes [5, 6]. The images are captured from every tilt angle within a range (typically *±*60^*°*^ with a 1^*°*^ interval), and the tilt series images are reconstructed into a 3D cryo-ET image, called a tomogram [6]. The tomograms are large grayscale volumes containing densities of multiple subcellular objects, including large membranes and smaller macromolecular complexes inside the cell [4, 7]. By analyzing these tomograms, it is possible to understand the morphology and organization of subcellular objects and link them to biological functions and mechanisms [8].

A crucial task for cryo-ET tomogram analysis is to segment the salient subcellular objects from the tomograms [7]. However, this task is significantly challenging for multiple reasons. First, the tomograms are extremely large 3D grayscale volumes with a typical size in the range of 4000 *×* 4000 *×* 2000 voxels [9, 7]. Even after binning four times across each axis, a tomogram is about 1000 *×* 1000 *×* 500 voxels. Segmenting such large volumes is inherently difficult. Moreover, cellular tomograms contain multiple subcellular objects of various shapes and sizes. A single tomogram can contain multiple large membranes and hundreds or even thousands of small macromolecular complexes. Segmenting such a large variety of objects across different scales is fundamentally challenging. Finally, cryo-ET tomograms are prone to artifacts and high noise due to the inherent properties of the sample and the imaging process [9, 6]. These artifacts and noise further complicate the segmentation process of cryo-ET tomograms.

In response to these challenges, several automated segmentation methods for cryo-electron tomograms (Cryo-ET) have been developed in recent years [7, 10, 11]. However, these methods have several drawbacks. Firstly, they can segment only one specific size of subcellular object-either more prominent membranes, or small macromolecular complexes. Second, many of them are based on supervised learning and require a large amount of annotated data to train. Annotating cryo-ET tomograms is a highly burdensome and time-consuming task. Consequently, the reliance on annotated data for training renders supervised segmentation models less effective and prone to human biases. Finally, cryo-ET tomograms of different cellular samples under various imaging conditions have significant domain differences, making the supervised segmentation model trained on one type of cellular tomogram ineffective for tomograms from a different domain. Although very recently, efforts have been made to develop methods with generalization ability across multiple domains of cryo-ET tomograms, their applicability is still limited [12]. Generalization across cellular cryo-ET tomograms of different organisms is highly challenging, given their high diversity and complexity.

As a solution to the problems mentioned above, we developed an unsupervised multiscale subcellular object segmentation approach for cellular cryo-electron tomograms, capable of segmenting both large structures, such as membranes, and small ones, like ribosomes, without the need for annotated data. Being an unsupervised method, our approach does not require users to annotate any tomogram from scratch to prepare training data, unlike supervised methods. Moreover, it can be independently applied to any cellular cryo-ET tomogram dataset, avoiding the difficulty of generalization across cellular tomograms from different experiments, organisms, and animals. Given a large cellular cryo-ET dataset, our framework uses only a few tomographic slices in a single tomogram for unsupervised training. Instead of relying on manual annotation, like supervised models, we leverage the foundational power of the Stable Diffusion model [13] to extract subcellular object features from these tomographic slices in an unsupervised manner. The approach we developed to this end is described in detail in the following sections.

We validated our unsupervised cryo-ET segmentation approach on two large publicly available cellular cryo-ET datasets-one containing 10 tomograms visualizing *S. Pombe* yeast cell sections, another with 100 tomograms visualizing *C. Elegans* worm cell sections. Both are cellular data sets of eukaryotic organisms that contain multiple subcellular objects of different sizes in a crowded cytoplasmic environment. We demonstrate that our method properly segments subcellular objects of diverse sizes and shapes, including large membranes and small macromolecular complexes, enabling multi-scale pattern mining. We observed that our unsupervised pipeline generates results comparable to those of supervised methods, which require exhaustive manual annotation of subcellular objects. These findings establish our approach as a robust and fully automated solution for multi-scale subcellular segmentation in large cryo-ET datasets, unlocking the potential to accelerate biological discovery by eliminating the bottleneck of manual annotation in cryo-ET analysis.

## Results

### Overview of our unsupervised segmentation pipeline

Our approach (Figure 1) aims to segment subcellular objects of diverse sizes and shapes—specifically macromolecules and membranes—from a collection of tomograms visualizing a cellular system, without relying on any manual annotations. Instead of requiring labels for the entire dataset, the approach only requires the user to select the training tomograms (which can be only one) and specify a small range of slices across the depth axis (referred to as slabs) within each selected tomogram. The selected slabs of the tomograms serve as the foundation of our unsupervised training pipeline. The slabs are large grayscale 2D images that are first enhanced using contrast adjustment techniques to highlight structural features. Each image is then split into four non-overlapping quarters to better capture localized details. These image quarters are used to extract high-level visual representations using a vision foundation model. Among the models evaluated, Stable Diffusion [13] was found to be particularly effective in capturing fine-grained and multi-scale features relevant for identifying different types of subcellular objects.

**Figure 1.**
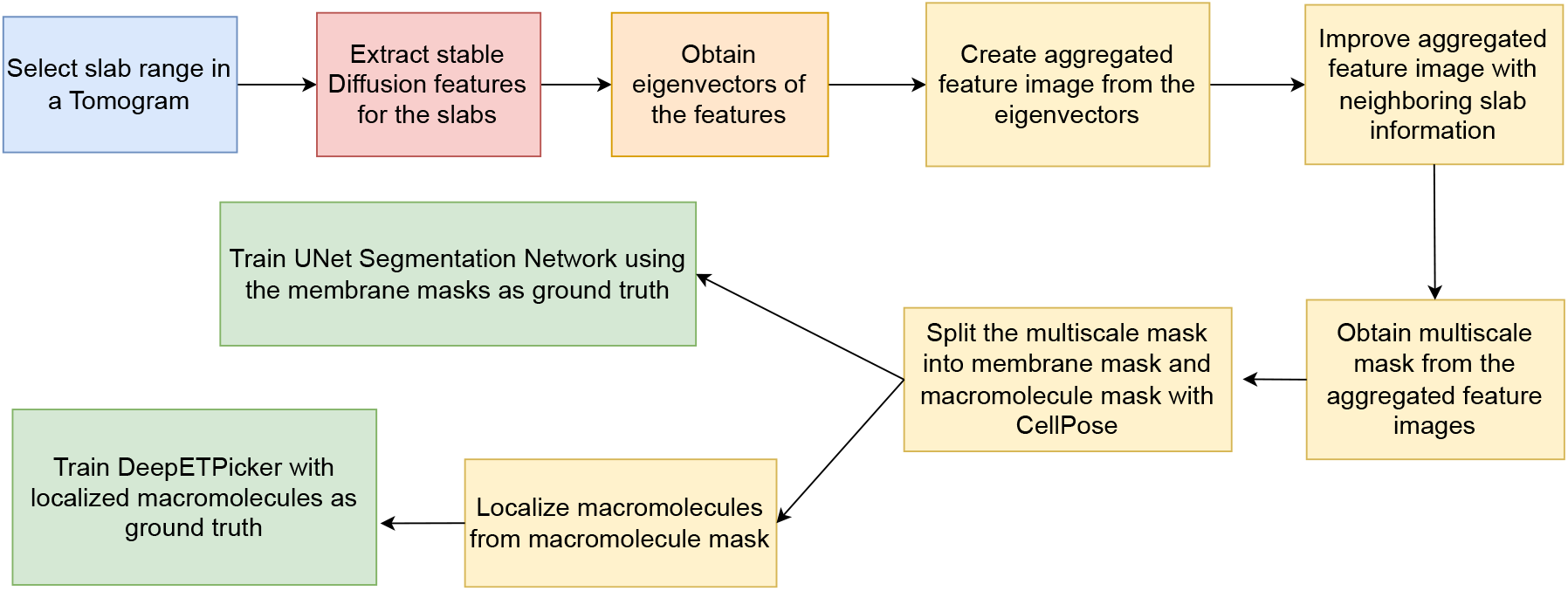
Our unsupervised segmentation pipeline workflow.

The visual representations obtained from the Stable Diffusion foundation model [13] are processed to capture spatial relationships between pixels using a mechanism inspired by how vision transformers analyze images [14]. The query-key vectors in the self-attention mechanism used in the UNet module of a Stable Diffusion foundation model [13] capture the essential features relevant for segmentation. Consequently, our method uses these query-key vectors to construct affinity graphs that reflect pixel-level similarity patterns across the image at different scales. To convert these patterns into meaningful object features, our method applies a spectral clustering-based technique that identifies groups of similar structures across different image patches. During the spectral clustering step, eigenvectors representing subcellular object features are derived from the affinity matrices. Among the eigenvectors, the most informative ones are selected based on how diverse and well-distributed their patterns are across the image. A custom heuristic is used to select and combine these features into a single, refined grayscale feature map. This map is then converted into a binary mask, which outlines regions in the image likely to contain subcellular objects. The binary mask highlights multiscale subcellular objects-both small macromolecules and larger membrane structures. Due to the sheer size and shape differences between macromolecules and membranes in the multiscale binary segmentation masks, they are split and processed differently in the downstream tasks. Our method utilizes a pre-trained deep learning model called CellPose [15] for detecting macromolecules from these multiscale binary masks. CellPose [15] is originally designed to segment cells and nuclei in fluorescence images. Since macromolecules in our aggregated binary mask resemble nuclei in fluorescence images, we found CellPose [15] to be effective at detecting macromolecule-like structures. After identifying and removing the macromolecule-like structures from the binary masks, morphological operations are used to isolate the remaining membrane regions. As a result, the method produces two distinct masks: one for macromolecules and one for membranes.

These masks are then used as pseudo-labels to train supervised models. A supervised UNet model [16] is trained on the membrane masks to predict membrane structures in all tomograms. Another learning-based weekly supervised method, DeepETPicker [11], is trained to detect macromolecule coordinates across the dataset. This pipeline—starting from minimal user input and ending with automated segmentation across all data—enables large-scale analysis of subcellular objects without manual annotations, using the inherent abilities of the Stable Diffusion foundation model [13].

### Unsupervised Segmentation of *S. Pombe* Cellular Tomograms

We first tested our unsupervised segmentation pipeline on cellular cryo-electron tomograms of *S. Pombe* cells [3]. We used 10 reconstructed tomograms directly from the CZI cryo-ET data portal. The tomograms were reconstructed from tilt-series movie frames, which were imaged by combining defocus and Volta potential phase plate (VPP). The images were four times binned, and the reconstructed tomograms had a pixel spacing of 13.48 Å. The tomograms contain large membranes, organelles, and small macromolecular complexes such as ribosomes and fatty acid synthase (FAS).

The tomograms had a zyx dimension of either 500 *×* 928 *×* 960 or 1000 *×* 928 *×* 960. We selected a random tomogram (named TS 0008) and visually inspected its slabs (xy slices across the z axis). We found slices *z* = 200 to *z* = 299 to have rich subcellular object features. We passed these 100 tomogram slices to our unsupervised pipeline. This accounts for only 1.33% of the total number of slices in the entire tomogram dataset. Using our unsupervised pipeline, we generated multiscale segmentation maps for these slices. We then split the multiscale segmentation maps into membrane masks and ribosome coordinates using CellPose [15], as described in the Methods section. The membrane masks were used as pseudo ground truth to train a supervised UNet segmentation network [16]. The macromolecule coordinates (88 obtained) and the selected tomogram TS 0008 were used to train a weakly-supervised particle picking approach, DeepETPicker [11]. Then the trained UNet was used to infer the membrane segmentation masks for all the slices in the tomogram, and the trained DeepETPicker [11] was used to infer macromolecule coordinates for the tomogram. We provided a step-by-step visualization for the entire process in Figure 2. Our segmentation models generalized well across all tomograms in the dataset. Then we used the trained UNet to infer membrane segmentation for all the remaining tomograms in the dataset. We present a few of our membrane segmentation results in Figure 3, along with the manual expert annotations as ground truth. To assess our performance in comparison to the supervised model, we trained the same UNet using manual expert annotations for the aforementioned slices of the selected tomogram as ground truth. We also provide the membrane segmentation results in Figure 3. The figure shows that our unsupervised method performs almost similarly to the supervised method and closely resembles the ground truth.

**Figure 2.**
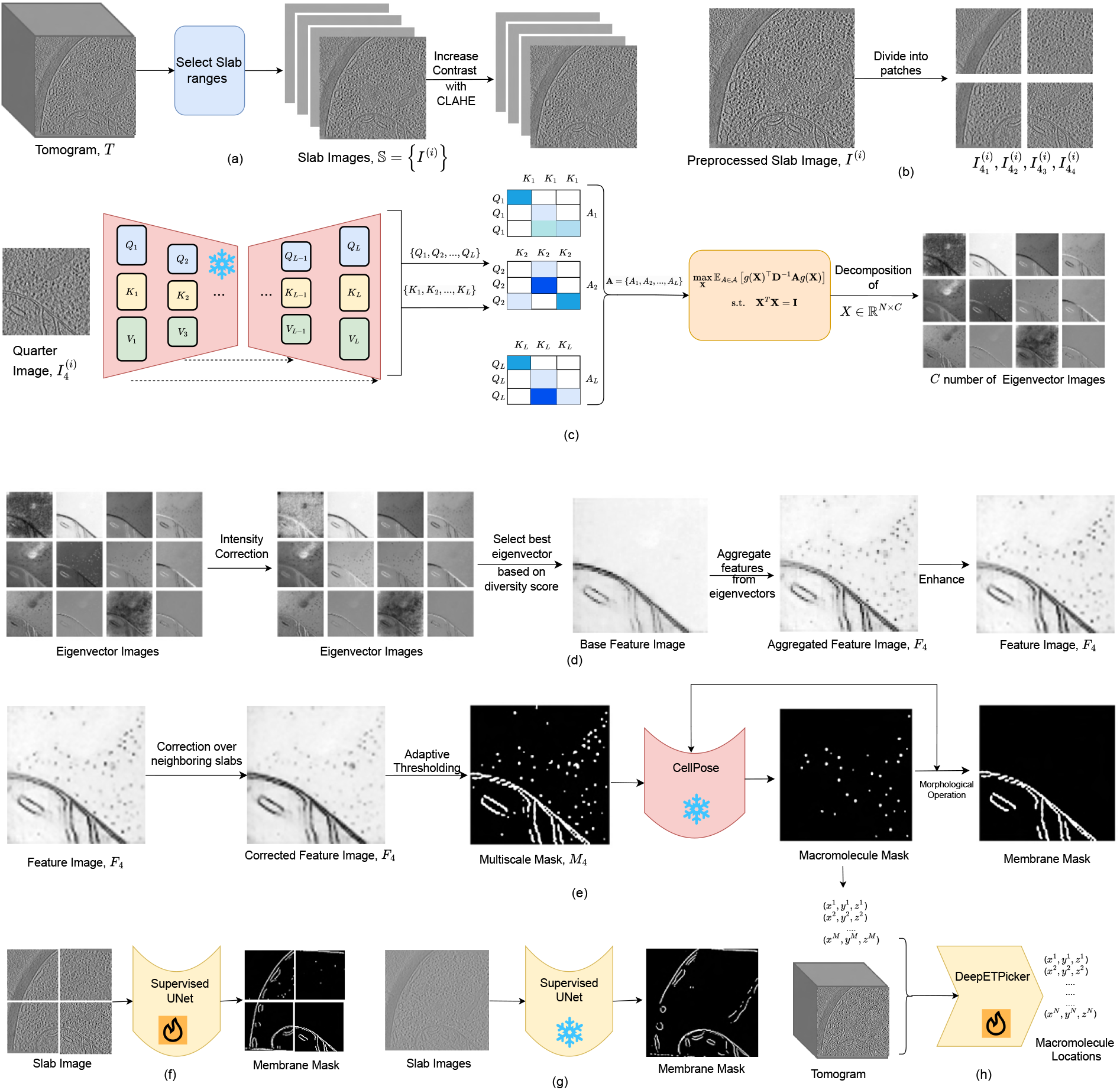
Our unsupervised segmentation pipeline with visual examples. (a) Selecting information-rich slabs from a tomogram and preprocessing the slabs. (b) Split the slabs into quarters. (c) Optimize eigenvector features of the quarter images from Stable Diffusion foundation model (d) Create feature image for the quarter images using the eigenvectors (e) Obtaining multiscale unsupervised segmentation mask from the feature image, spliting the multiscale mask to membrane and macromolecule masks (f) train a supervised UNet with predicted unsupervised membrane masks as ground truth (g) Use the trained UNet to infer membrane masks for other slabs in the tomogram. (h) Train a DeepETPicker model with the locations extracted from the predicted unsupervised macromolecule mask and obtain tomogram-wide locations of macromolecules.

**Figure 3.**
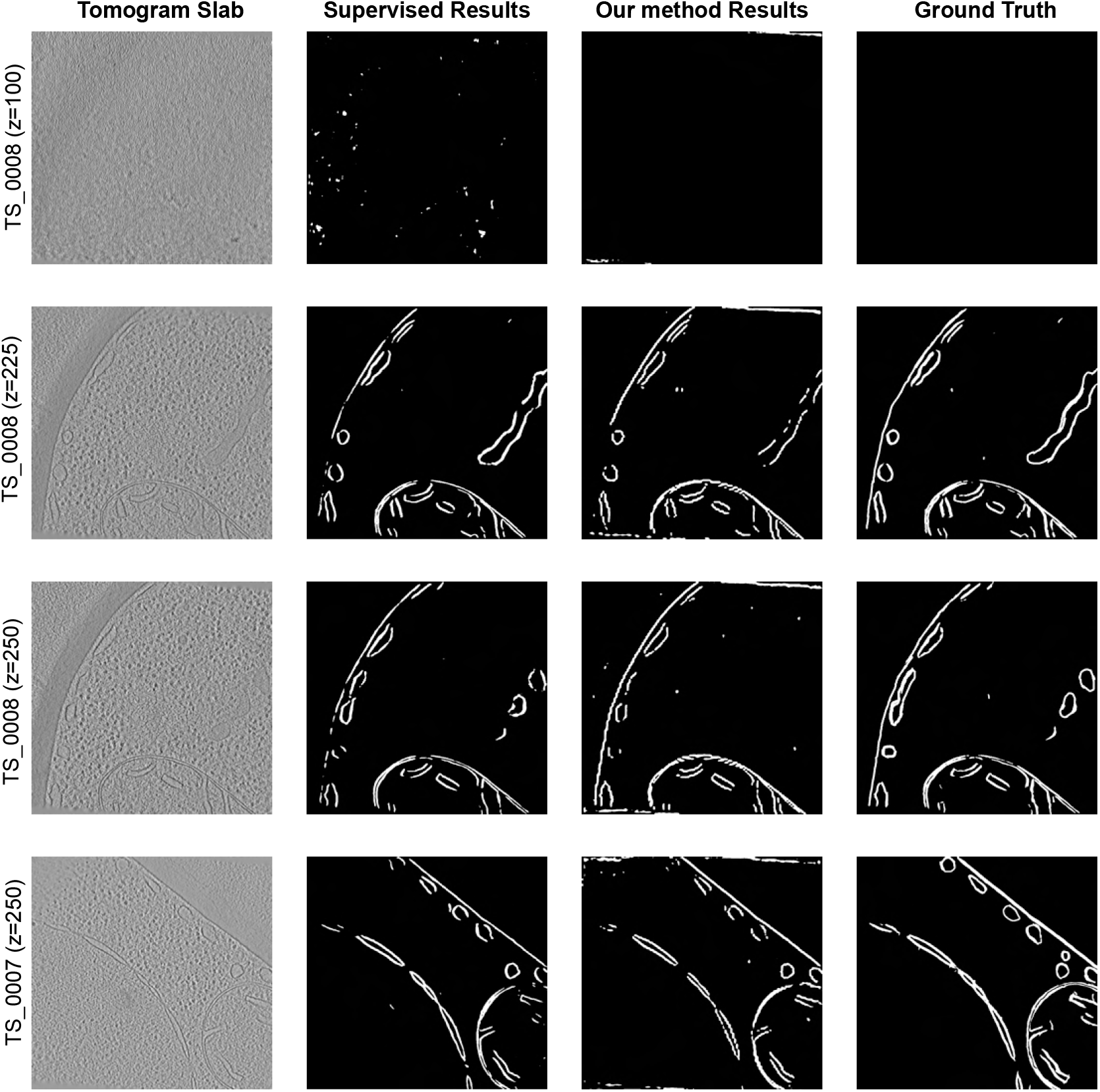
Membrane segmentation results on *S. Pombe* Tomograms. The image shows several slabs of training tomogram (TS 0008) and another tomogram with the expert annotated membrane mask as ground truth, and the predicted mask by supervised UNet and our unsupervised approach.

We further quantitatively evaluated the performance of our method and the supervised version for membrane segmentation. We calculated the Dice Coefficient (as defined in the supplementary document) for both methods with respect to the expert-annotated ground truths. The quantitative result is provided in Table 1. It can be observed that the supervised version performs only 4% (23% for training tomogram TS 0008) higher than our unsupervised method. This suggests that our unsupervised method achieves competitive performance despite the absence of manual labels during training. The relatively small performance gap highlights the effectiveness of our approach in capturing membrane structures directly from the data.

**Table 1.** Dice score comparison between two training methods for membrane segmentation on VPP *S. Pombe* cellular cryo-ET datasets. The higher score indicates better segmentation.

| Model Type | TS_0001 | TS_0002 | TS_0003 | TS_0004 | TS_0005 | TS_0006 | TS_0007 | TS_0008 | TS_0009 | TS_0010 | Overall |
| --- | --- | --- | --- | --- | --- | --- | --- | --- | --- | --- | --- |
| **Supervised** | 0.045 | 0.323 | 0.439 | 0.323 | 0.366 | 0.263 | 0.354 | 0.665 | 0.348 | 0.113 | 0.324 |
| **Unsupervised** | 0.06 | 0.379 | 0.545 | 0.394 | 0.234 | 0.248 | 0.356 | 0.508 | 0.241 | 0.123 | 0.309 |

We further used the trained DeepETPicker [11] to infer macromolecule coordinates for all the remaining tomograms in the dataset. We quantitatively evaluated our macromolecule localization results and compared their performance with that of CrYOLO[17] and DeepETPicker [11], which were trained with varying numbers of ground truth coordinates. Similar to recent particle picking methods, we evaluated the performance of our method and the supervised methods by calculating the F1 scores (definition in the supplementary document). We present the quantitative results in Table 2. It can be observed that our approach, which is essentially a DeepETPicker model trained with 88 pseudo-ground truth macro- molecule coordinates predicted by the unsupervised method, performs significantly better (+22.84%) than the DeepETPicker model trained with 100 ground truth coordinates. It even performs significantly better (38.71%) than the supervised CrYOLO method trained with 500 ground-truth macromolecule coordinates. However, the performance of supervised DeepETPicker [11] trained with 500 ground truth macromolecule coordinates performed much better (+32.55%) than our unsupervised approach, suggesting that processing more slabs for the unsupervised pipeline may yield better macromolecule localization performance.

**Table 2.** F1 Scores of different methods on VPP *S. Pombe* cellular cryo-ET datasets. The higher score is better. The value of *n* in supervised models represent the number of training particle coordinates.

| Model Type | Model Name | TS.0001 | TS.0002 | TS.0003 | TS.0004 | TS.0005 | TS.0006 | TS.0007 | TS.0008 | TS.0009 | TS.0010 | Overall |
| --- | --- | --- | --- | --- | --- | --- | --- | --- | --- | --- | --- | --- |
| Supervised | CrYOLO (n=10) | 0.01 | 0.01 | 0.01 | 0.00 | 0.01 | 0.01 | 0.00 | 0.00 | 0.00 | 0.00 | 0.01 |
|  | CrYOLO (n=100) | 0.37 | 0.28 | 0.29 | 0.29 | 0.24 | 0.13 | 0.07 | 0.19 | 0.26 | 0.38 | 0.25 |
|  | CrYOLO (n=500) | 0.36 | 0.18 | 0.44 | 0.28 | 0.29 | 0.25 | 0.12 | 0.42 | 0.31 | 0.42 | 0.31 |
|  | DeepETPicker (n=10) | 0.23 | 0.19 | 0.17 | 0.13 | 0.27 | 0.14 | 0.11 | 0.11 | 0.21 | 0.16 | 0.17 |
|  | DeepETPicker (n=100) | 0.70 | 0.23 | 0.70 | 0.05 | 0.52 | 0.41 | 0.45 | 0.13 | 0.37 | 0.44 | 0.35 |
|  | DeepETPicker (n=500) | 0.77 | 0.59 | 0.66 | 0.23 | 0.72 | 0.65 | 0.49 | 0.22 | 0.67 | 0.67 | 0.57 |
| Unsupervised | Our Method | 0.42 | 0.45 | 0.41 | 0.35 | 0.59 | 0.43 | 0.28 | 0.44 | 0.48 | 0.43 | 0.43 |

We also provide visualizations for a few of our macromolecule localization results in Figure 4. Figure 4 visualizes macromolecule locations predicted by DeepETPicker trained with 100 ground truth coordinates, and our unsupervised pipeline together with the expert annotations as ground truth. It can be observed that our unsupervised pipeline more closely resembles the ground truth compared to the supervised alternative in the figure, supporting the higher F1 score achieved by our pipeline.

**Figure 4.**
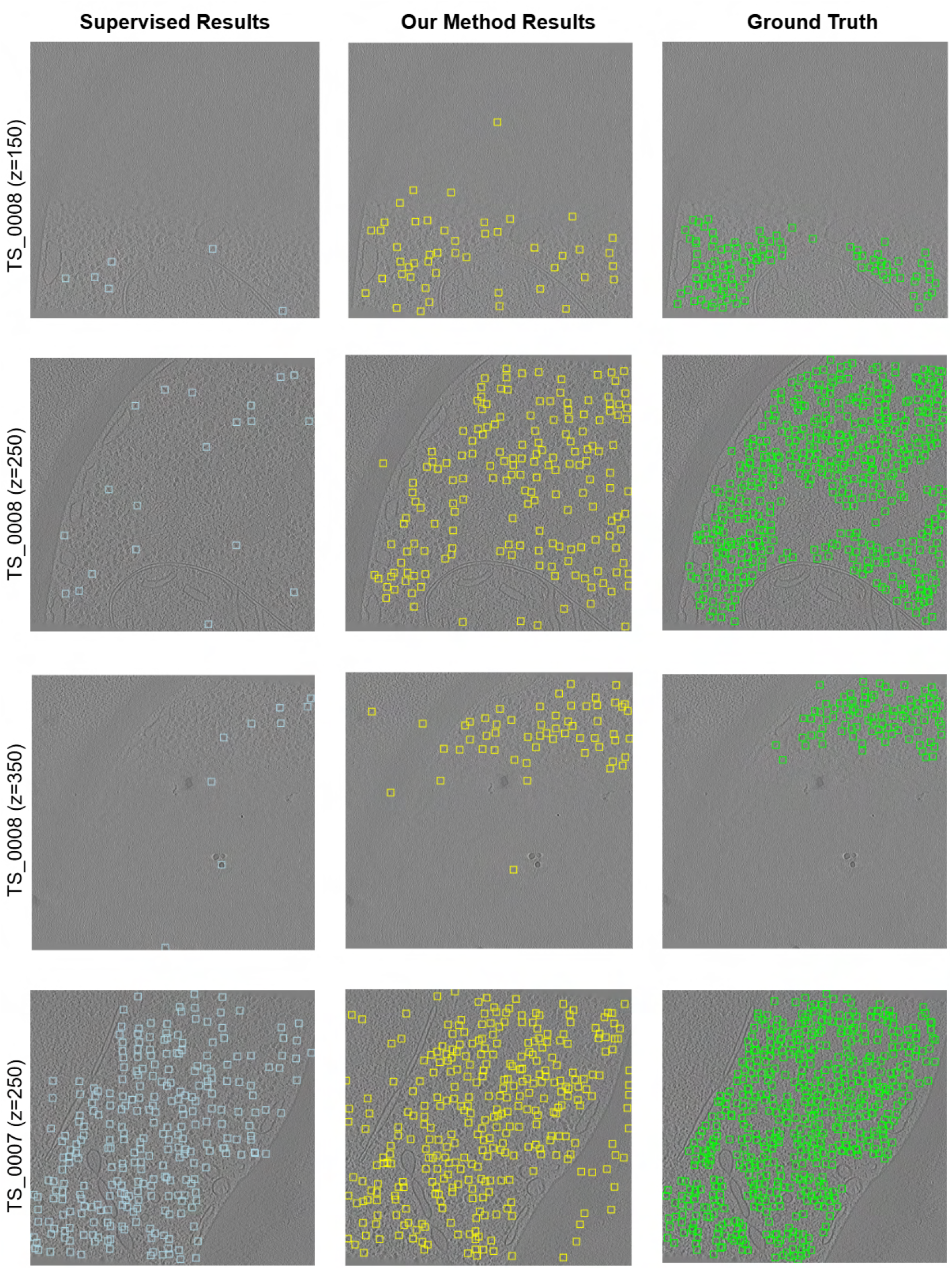
Macromolecule localization results on *S. Pombe* Tomograms. The image shows several slabs of training tomogram (TS 0008) and another tomogram with annotated macromolecules drawn as bounding boxes. The expert annotated macromolecule locations (green boxes) are provided as ground truth. The locations predicted by a supervised DeepETPicker [11] trained with 100 ground truth coordinates are provided as supervised results with cyan boxes. The locations predicted by our unsupervised pipeline are provided with yellow boxes.

We also evaluated the deep metric learning based model TomoTwin [12] on the training tomogram TS 0008. TomoTwin requires a single reference particle and outputs a list of predicted particle locations based on the similarity and dissimilarity of embeddings extracted from subvolumes in the tomogram. With TomoTwin, we achieved an F1 score of only 0.027. The performance drop is due to the fact that TomoTwin was trained on simulated single particle tomograms that do not well represent the complexity of the cellular tomogram used in our experiment. This also highlights the challenge of building a generalized model with cross-dataset generalization, due to significant domain differences in cellular cryo-ET images.

### Unsupervised Segmentation of *C. Elegans* Cellular Tomograms

After evaluating and validating our pipeline on *S. Pombe* tomograms, we applied our segmentation pipeline on a large dataset of *C. Elegans* cellular cryo-electron tomograms. The dataset contains 100 tomograms, visualizing various cell sections of *C. Elegans* cells. The tomograms have a uniform voxel spacing of 7.56 Å. These tomograms contain multiple subcellular objects, including membranes, ribosomes, and other cellular components.

The tomograms in the dataset were of uniform size with a zyx dimension of 500 *×* 1024 *×* 1024. Since no Volta phase plate was employed during image acquisition, the resulting tomograms exhibited a higher noise level compared to the *S. Pombe* dataset. To mitigate this and enhance feature extraction, we applied the CCP-EM denoiser [18] prior to segmentation. Then, we selected a random tomogram (named Position 508 3) and visually inspected its slabs (xy slices across the z axis). We found slices *z* = 200 to *z* = 349 to have rich subcellular object features. We passed these 150 tomogram slices to our unsupervised pipeline. This accounts for only 0.47% of the total number of slices in the entire tomogram dataset. Similar to our experiment with *S. Pombe* tomograms, we obtained a trained UNet [16] for membrane segmentation and trained DeepETPicker for macromolecule localization. We used these deep networks to infer results for all the remaining tomograms and their slices in the dataset.

Unlike the *S. Pombe* tomograms, the *C. Elegans* tomograms lack well-defined expert-annotated ground truths, which precludes direct quantitative evaluation of segmentation performance. Despite this limitation, we conducted a comprehensive qualitative assessment of the results. The segmentation outputs exhibit consistency with expected biological morphology and structural boundaries, suggesting that our method captures biologically meaningful features even in the absence of annotations. We showcase several examples of multiscale segmentation in Figure 5, where diverse subcellular objects—ranging from large membrane systems to smaller macromolecular assemblies—are detected. These results were generated in a fully automated, data-driven manner without any human supervision or manual annotation.

**Figure 5.**
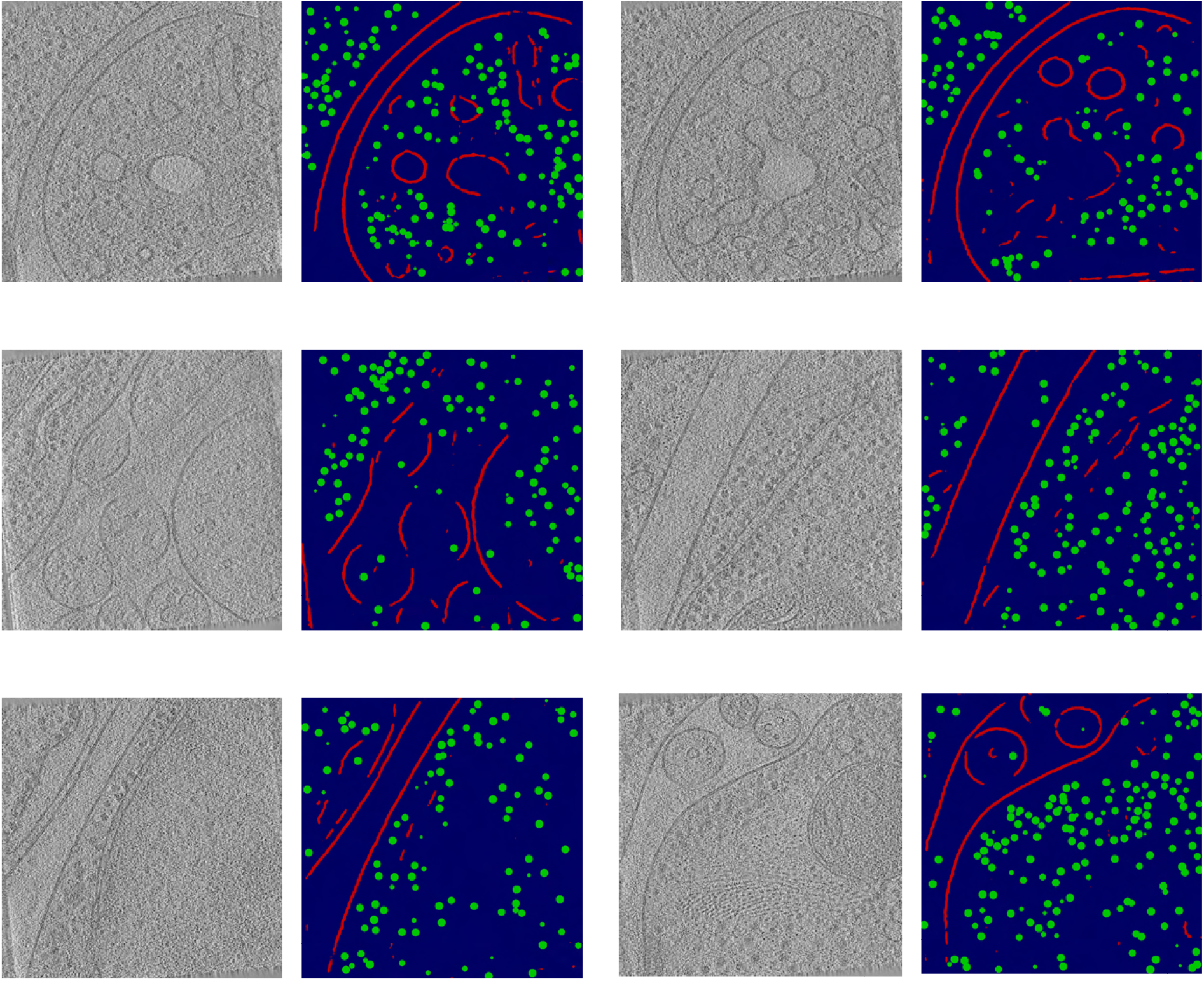
Multiscale segmentation results on *C. Elegans* Tomograms. The image shows pairs of cryo-ET slabs and their corresponding predictions. In the predicted mask, the red lines indicate membranes and the green dots represent macromolecules.

## Discussion

Contemporary structural biology bridges the fields of structural and cell biology, encompassing multiple scales of subcellular objects. Cellular cryo-electron tomography is a powerful technology that directly visualizes subcellular objects of diverse sizes and shapes. Recent technological advancements have led to a rapid increase in the availability of cellular cryo-ET datasets. Automated segmentation of subcellular objects from these datasets is essential for advancing our understanding of cellular architecture. While efforts have been made to develop automated methods, most are currently limited to objects with specific shapes or sizes and rely heavily on supervised learning, which in turn requires extensive manual annotation. To address these limitations, we developed a learning-based, unsupervised multi-scale subcellular object segmentation approach in this work, leveraging the power of AI, particularly a vision foundation model. The method is effective in annotating a cellular cryo-electron tomogram almost independently, significantly reducing the need for manual intervention.

However, our unsupervised segmentation pipeline primarily segments the membranes and macro-molecules inside the cell, not individual organelles. For example, it segments the outer membrane of the Endoplasmic Reticulum (ER), not the region enclosed by the ER membrane. This is obvious given the nature of the cryoelectron tomograms, where the objects are identified with alternations in contrast. Consequently, to segment individual subcellular object regions without any contrast change, unsupervised methods are not suitable, and a form of bias through supervised annotation is compulsory. Nevertheless, individual subcellular objects, such as mitochondria, ER, Golgi apparatus, etc., can already be distinguished through the segmentation of their characteristically shaped membranes [4]. As for macromolecules, our unsupervised pipeline localized large complexes such as ribosomes and fatty acid synthases. Given their size, these macromolecules are generally visible through contrast changes in the cryo-electron tomograms. For smaller and rarer macromolecules in cellular tomograms, manual picking is still necessary.

In our experiments, we used only a few (100 for *S. Pombe* and 150 for *C. Elegans*) slabs from a single tomogram to train our unsupervised pipeline. However, the performance can be largely improved by selecting more slabs across multiple tomograms. In our unsupervised pipeline, the only computationally expensive step is the eigenvector optimization process, which takes around 7*−*8 minutes on a single NVIDIA A5000 GPU for a single slab. One drawback is that this processing time scales linearly with the number of slabs, resulting in a significant increase in processing time for a large number of slabs. However, this computational efficiency issue can be solved with multi-GPU parallelization and batch processing in the eigenvector learning process.

Our method complements existing methods, such as Membrain-Seg [10, 19], for membrane segmentation in cellular cryo-ET tomograms. Whereas Membrain-Seg is a generalized model trained on multiple manually annotated cryo-ET tomograms, it may still generate spurious segmentations on new tomograms, particularly those having much differently shaped membranes compared to their training data. Being an unsupervised approach, our method would not face training data bias. In practice, the membrane segmentation provided by our method and by Membrain-seg [10] can be used together. Furthermore, the membrane segmentation produced by our method can be imported into ColabSeg [20] and edited for ground truth preparation, offering a far more efficient alternative to manual annotation from scratch.

For macromolecule localization, our method is designed to detect macromolecules of various types, though it may also produce some false positives. However, these false positives can often be filtered out during downstream 3D classification. For example, the localized subtomograms may include a mixture of ribosomes, FAS, fiducials, and noise from cellular tomograms. These can subsequently be classified into distinct structural classes using tools such as RELION [21] or DISCA [9]. Additionally, providing a larger number of input slabs during localization can help increase the number of true positive macromolecules while reducing the presence of noise particles or false positives.

In summary, our method introduces a pioneering, fully unsupervised learning-based approach for segmenting multiple subcellular objects of varying sizes and shapes, including membranes and macromolecules, directly from cellular cryo-ET tomograms. By automatically generating membrane masks and localizing potential macromolecular components, it offers an effective starting point for biologists seeking to explore complex and previously uncharacterized cellular systems with cryo-ET. Beyond its biological utility, our framework showcases the capability of Stable Diffusion foundation models for 3D segmentation of large, complex, and hierarchically organized volumetric data, establishing a versatile paradigm of broad relevance to the computer science, machine learning, computational biology, and medical imaging communities.

## Methods

### Preprocessing

Our pipeline takes a set of *K* slab images, 𝕊 = {*I*^(*i*)^ ∈ ℛ^*H×W*^, *i* ∈ [*K*]}, from the selected tomograms as input. Initially, our pipeline applies multiple image processing steps to these images to enhance their contrast. Among the processing steps, we first apply contrast stretching, where the pixel intensity values are normalized to a fixed range, ensuring better visibility of the objects and bringing the values into the expected range for images. To further improve local contrast, Contrast Limited Adaptive Histogram Equalization (CLAHE) [22] is applied. CLAHE divides the image into small tiles and redistributes the intensity values within each tile while limiting the amplification of noise. This step enhances subtle features that may otherwise be obscured during global normalization. We use the opencv package to apply CLAHE to the input images. We then divide each preprocessed image *I*^(*i*)^ ∈ 𝕊 into 4 equally sized nonoverlapping quarters 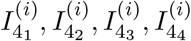. We refer to the set of all the quarter images extracted from all *I*^(*i*)^, *i* ∈ [*D*] as 𝕊_4_.

### Feature processing from Stable Diffusion Foundation Model

For each of the images 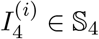, we extract the feature representations from a Stable Diffusion foundation model [13] directly from [23] without any training or fine-tuning. These features are derived from the attention layers of the conditional UNet module of the Stable Diffusion model [13]. A brief discussion of the Stable Diffusion model and attention mechanism is provided in the supplementary document. During feature extraction, we consider all attention layers of the conditional UNet module in the Stable Diffusion model, rather than focusing solely on the last layer, as several feature extraction methods do [24, 25].

The input image quarters 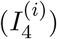 in this step (typically dimension of 512 *×* 512) are split into patches and then flattened and used as visual tokens. In our experiments, we use a patch size of 8 and consequently obtained 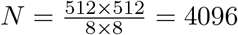 visual tokens. We use the attention layers of the UNet module of the Stable Diffusion foundation model to extract query and key embeddings ((*Q*_1_, *K*_1_), (*Q*_2_, *K*_2_), …, (*Q*_*L*_, *K*_*L*_), where *L*=Number of attention layers) and (*Q*_*l*_, *K*_*l*_) are (query, key) embedding pairs of *l*-th layer. Both *Q*_*l*_ and *K*_*l*_ have a dimension of *N ×* 1, where *N* is the number of visual tokens. We use query-key embedding pairs since query-key embedding pairs in vision transformers provide critical spatial-level information for segmentation tasks [14]. Using these embeddings, we obtain a set of affinity matrices **A** = {*A*_1_, *A*_2_, …, *A*_*L*_}. Each affinity matrix *A*_*l*_ ∈ ℝ^*N ×N*^, *l* ∈ [*L*] is computed as:

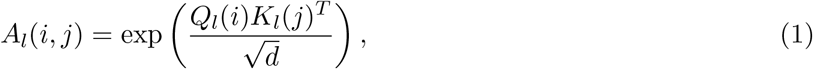

Here *A*_*l*_(*i, j*) is the affinity between data samples *i* and *j* in the affinity matrix *A*_*l*_. *d* is the feature dimension. *Q*_*l*_(*i*) is query for *i*-th data point and *K*_*l*_ is the key for *j*-th data point in the *l*-th attention layer. In our Stable Diffusion foundation model [23], *N* = 4096 and *L* = 16.

To extract meaningful subcellular object features from **A**, we followed a spectral clustering approach [14]. In this approach, an eigenvector matrix with *C* eigenvectors, *X* ∈ ℝ^*N ×C*^ is derived from *A* by solving the following eigenvalue problem.

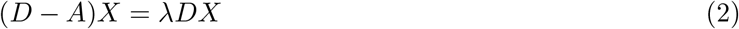

where, *D* ∈ ℝ^*N ×N*^ is the diagonal degree matrix of *A. λ* is the eigenvalues and *X* is the eigenvector. The objectness features are typically distributed across all attention layers. Consequently, to get rich objectness features, we use all layers of attention. Instead of a single *A*, we use **A** = {*A*_1_, *A*_2_, …, *A*_*L*_} and to account for the increasing |**A**|, we solve the following approximation of Eqn 2.

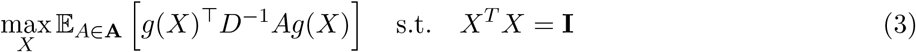

Here, *g*(.) is a function that takes *X* as input and returns a matrix matching the size of *A*. For further details on the derivation of the approximation, we refer to [14].

We parametrized the eigenvector *X* as a learnable feature map and optimized the objective function in Eqn. 4 with gradient-based optimization. ∥.∥_*F*_ in Eqn. 4 denotes Frobenious norm.

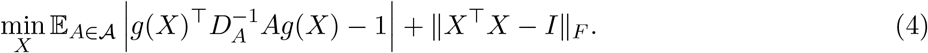

The optimal *X* is obtained through this optimization. *X* contains *C* eigenvectors each with dimension *N ×* 1. We utilize the PyTorch autograd framework to implement this optimization, which is similar to the process of training neural networks.

To separate distinct features across different eigenvectors in *X*, we enforce an orthogonal representation across different channels of *X*. To this end, similar to [14], we find the eigenvectors *U* of a matrix *X*^*T*^ *X* ∈ ℝ^*C×C*^ and then update *X* using *X* = *XU*.

### Converting eigenvectors to segmentation mask with adaptive thresholding

For each image in ℝ_4_, we obtain *C* number of eigenvectors with dimensions *N ×*1. In our case, *N* = 4096. We reshape the eigenvectors as grayscale eigenvector images of size 64 *×* 64. We introduced a new heuristic, termed diversity score, which measures the feature diversity across a grayscale image. We define it as follows:

**Diversity score**

Let *I* ∈ ℝ^*H×W*^ be a grayscale image. Let *p* denote the patch size, *o* the overlap, and *s* = *p − o* the stride. Define the set of extracted patches as:

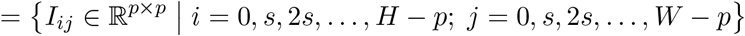

For each patch *I*_*ij*_ ∈ 𝒫, we compute the standard deviation of its pixel intensities: *σ*_*ij*_ = std(*I*_*ij*_) Now, let the set of these local standard deviations be: ***σ*** = {*σ*_*ij*_ | *I*_*ij*_ ∈ 𝒫}

Then, the **diversity score** of the image is defined as: DiversityScore(*I*) = std(***σ***)

We calculated the diversity score for each eigenvector image and selected the one with the highest score as our primary segmentation feature image, which we referred to as the ‘base’ image. We iteratively updated this ‘base’ image by merging additional eigenimages to the base image where the following three conditions hold:

- The eigenimage has a high similarity with the base image
- Adding the eigenimage to the base image increases the diversity score of the base image
- Adding the eigenimage to the base image does not result in more noise (measured by very high contour count)

We kept updating the ‘base’ image until none of the above conditions were met. We regarded the updated ‘base’ image as the ‘aggregated feature’ image. Thus, for each image 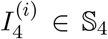, we obtain an aggregated feature image 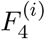. Given that the shape of the eigenvector images was 64 *×* 64, the shape of each aggregated feature image is also 64 *×* 64. We reshape and interpolate the feature image 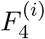 to upsample it back to the size of 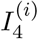 using opencv.resize.

Afterwards, we corrected the upsampled aggregated feature images based on the feature images from their neighboring slabs across the depth (z) axis. Since the neighboring slabs across the z-axis visualize the subcellular objects at almost similar spatial positions, the aggregated feature images for them should be highly similar. Consequently, we perform the following two actions:

Assume, 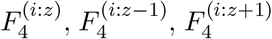 is the aggregated feature image obtained for quarter image 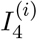 at *z*-th, (*z −* 1)-th, and (*z* + 1)-th slab respectively.

- If 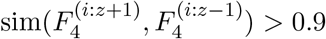 and 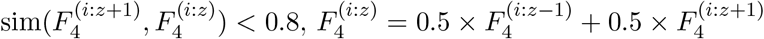.
- Otherwise, 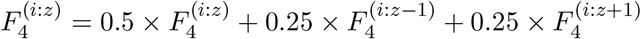

The first condition deals with the erroneous feature images. If a feature image has low similarity with its z-axis neighbors, whereas the neighbors have high similarity between them, then the feature image is probably erroneous. In such a case, we fix the feature image by setting it to be the average of the corresponding feature images in the neighboring slices. To calculate similarity between two images, we use the Structural Similarity Index Measure (SSIM) (discussed in the supplementary document).

The second condition takes into account the continuity across the z-axis. Although we regard the slabs as independent 2D images, in reality, the neighboring slabs are interdependent. The second condition thus updates the aggregated feature image by taking into account the contents of the feature images from the adjacent slabs.

After correcting the aggregated feature images based on neighboring slab information, we segmented the subcellular objects from this corrected aggregated feature image using the following procedure.

We first applied Gaussian smoothing to the ‘aggregated feature’ image. We then performed Gaussian adaptive thresholding on the smoothed image to convert it from grayscale to a binary image. In adaptive thresholding, for each pixel in the ‘aggregated feature’ image, we calculated a threshold based on the pixel values in its local neighborhood and a constant *C*. The local neighborhood is defined by a block size *b*. In our experiments, we used *b* = 15 and *C* = 3. If the pixel value is greater than the threshold, it is regarded as foreground (value 255); otherwise, it is considered background (value 0). Further details on the adaptive thresholding algorithm are provided in the supplementary document.

After adaptive thresholding, for each image 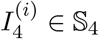, we obtain a binary mask 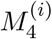.

### Determining masks for membranes and macromolecules with pretrained CellPose

The binary mask obtained through adaptive thresholding typically includes both membranes and macro-molecules. However, these structures differ significantly: membranes form long, continuous shapes, whereas macromolecules appear as small, nearly spherical objects. To handle these differences, the binary mask is separated into two distinct components—one for membranes and one for macromolecules. The pre-trained CellPose model [15], a deep learning tool initially designed for cell segmentation in fluorescence microscopy, is used for the decomposition. Since macromolecules in our binary mask appear similar to cells in such images, CellPose [15] is used to extract their masks. The remaining foreground regions, primarily representing membranes, are refined using morphological operations such as closing and dilation.

### inference of membrane segmentation masks

To infer membrane masks for the remaining slices in the tomogram in the dataset, a UNet model [16] is trained from scratch. The UNet model [16] takes a slab image *I* as input and predicts a binary segmentation mask. The training of UNet is performed over 30,000 iterations using a batch size of 8. Optimization is performed using stochastic gradient descent (SGD) with an initial learning rate of 0.1, momentum of 0.9, and a weight decay of 1 *×* 10^*−*5^ to prevent overfitting. A StepLR learning rate scheduler is applied to decay the learning rate by a factor of 0.1 every 10,000 iterations. To ensure reproducibility, all experiments are conducted with a fixed random seed. The loss function combines Dice loss and cross-entropy loss to capture both region-level and pixel-level segmentation accuracy effectively. Data augmentation techniques, such as random flipping and random rotation, are employed to enhance model generalization. The Dice coefficient on the validation set is used as the primary evaluation metric to monitor segmentation performance.

### Inference of macromolecule coordinates

Our multiscale segmentation mask provides only a few particles in the selected slabs of the tomogram. To infer the remaining particles, we train a weakly supervised learning-based particle picking network, termed DeepETPicker [11], using the multiscale mask and macromolecule coordinates as ground truth. To this end, we use the default settings provided in the DeepETPicker code [26]. Further details on DeepETPicker training and inference are provided in the supplementary document.

## Data Availability

The tomograms used for segmentation in this work are all publicly available at the CZI cryo-ET data portal. The *S. Pombe* tomograms are available on CZI cryo-ET data portal with dataset ID: DS-10001 and also as EMPIAR-10988. The *C. Elegans* tomograms are available on CZI cryo-ET data portal with dataset ID: DS-10004.

## Code Availability

The code to run our method, along with the necessary instructions, will be publicly available upon publication.

## Acknowledgement

This work was supported in part by U.S. NSF grant DBI-2238093.

## Author Contributions

M.R.U. and M.X. conceived the research. M.R.U. designed and implemented the unsupervised segmentation pipeline. T.N. implemented the UNet-based membrane segmentation module. K.G. developed the CellPose-based mask splitting module. M.R.U., T.N., H.M.S.T., and K.G. performed the experiments.

M.R.U. and H.M.S.T. drafted the manuscript. All authors reviewed and edited the manuscript.

