## Supplementary document for "Unsupervised Multi-scale Segmentation of Cellular cryo-electron Tomograms with Stable Diffusion Foundation Model"

#### 1 Methods

##### 1.1 Contrast Limited Adaptive Histogram Equalization (CLAHE)

Contrast Limited Adaptive Histogram Equalization (CLAHE) is an advanced image processing technique designed to enhance local contrast in grayscale images, making subtle details and structures more visible without excessively amplifying noise. Unlike traditional histogram equalization methods, which apply global intensity transformations across the entire image, CLAHE performs adaptive equalization locally within small image regions, thereby preserving local contextual information and mitigating global artifacts.

CLAHE operates through the following primary steps:

1. **Division into Tiles:** The input grayscale image is partitioned into multiple non-overlapping, small regions or tiles (typically of equal size). This localized processing approach enables adaptive adjustment to spatially varying illumination and texture across the image.

2. **Local Histogram Computation:** For each tile, a histogram of pixel intensities is computed, capturing the local distribution of intensity values.

3. **Histogram Clipping:** To avoid excessive noise amplification, CLAHE introduces a clipping parameter, commonly referred to as the *clip limit*. Any histogram bin exceeding this predefined limit is clipped, and the excess pixel counts are redistributed uniformly across the entire histogram. This clipping process effectively reduces unwanted noise amplification in homogeneous regions.

4. **Histogram Equalization:** After clipping, each histogram is transformed into a cumulative distribution function (CDF), which is used to remap the intensity values within the corresponding local region. Pixels within each tile are then equalized according to their local CDF, resulting in an enhancement of local contrast and improved visibility of subtle image features.

5. **Interpolation between Tiles:** To achieve smooth intensity transitions and eliminate tile-edge artifacts, bilinear interpolation is typically applied across the boundaries of adjacent tiles, resulting in a seamless, globally consistent enhancement of the entire image.

###### Mathematical Formulation of CLAHE

Given an image region (tile) with intensity levels ranging from 0 to $L-1$, CLAHE computes the clipped local histogram $h(i)$, where $i$ denotes the intensity level. The histogram clipping process is expressed as:

$$h_{\text{clipped}}(i) = \begin{cases} h(i), & h(i) \leq C_{\text{clip}} \\ C_{\text{clip}}, & h(i) > C_{\text{clip}} \end{cases}$$

where $C_{\text{clip}}$ is the predefined clip limit. The excess histogram counts, defined as $\sum_i [h(i) - C_{\text{clip}}]^+$, are then redistributed uniformly across all histogram bins. Following this redistribution, the cumulative distribution function (CDF) is computed as:

$$\text{CDF}(i) = \frac{(L-1)}{N} \sum_{j=0}^{i} h_{\text{clipped}}(j),$$

where $N$ is the total number of pixels in the local region. Finally, each pixel intensity $I(x, y)$ within the tile is mapped to its enhanced value $I'(x, y)$ using:

$$I'(x, y) = \text{CDF}(I(x, y)).$$

In our segmentation pipeline, CLAHE plays a crucial role in preprocessing. By adaptively enhancing local contrast, it significantly improves the visibility of subtle biological structures in cryo-electron tomography (Cryo-ET) images, facilitating more accurate downstream unsupervised segmentation. Specifically, CLAHE helps in highlighting membrane boundaries and fine macromolecular details, which would otherwise remain obscured due to low inherent contrast and uneven illumination typically present in Cryo-ET datasets.

We implement CLAHE using standard image processing libraries (such as OpenCV), with carefully selected parameters (e.g., tile size and clip limit) optimized empirically for our Cryo-ET data, thus ensuring effective noise suppression while preserving essential biological structures.

#### 1.2 Vision Foundation Models

Vision Foundation Models (VFMs) are large-scale, pre-trained neural networks designed to perform a wide range of computer vision tasks with minimal task-specific supervision. Inspired by foundation models in natural language processing (like GPT), VFMs are trained on massive, diverse image datasets using self-supervised or weakly supervised learning. They learn generalized visual representations that can be adapted or fine-tuned for downstream tasks such as classification, segmentation, object detection, and image captioning.

Popular examples include CLIP (Contrastive Language–Image Pre-training), DINO (self-distillation with no labels), SAM (Segment Anything Model), Point-level Region Contrast, and Stable Diffusion. VFMs often rely on architectures like Vision Transformers (ViT) and use techniques such as contrastive learning, masked image modeling, or image-text alignment.

In this work, we use VFM as a feature extractor for cryo-ET image slices. In Figure 1, we show extracted features for a sample cryo-ET image slice for multiple VFMs. We found that the stable diffusion model provided us with the best features overall, a similar phenomenon observed by [**?**]. Consequently, we used stable diffusion as a foundational model in our unsupervised segmentation pipeline.

#### 1.3 Stable Diffusion Foundation Model

The Stable Diffusion model is a state-of-the-art latent text-to-image generative model that excels at generating photorealistic images conditioned on textual descriptions. Developed as a diffusion-based generative framework, Stable Diffusion integrates powerful deep architectures, including a Variational Autoencoder (VAE), a UNet denoising backbone incorporating a Vision Transformer (ViT), and a Transformer-based text encoder. This sophisticated architecture enables robust multimodal representation learning.

Unlike conventional generative models, Stable Diffusion operates within a learned latent space rather than directly at pixel resolution, significantly reducing computational complexity while preserving semantic fidelity. It employs a **Denoising Diffusion Probabilistic Model (DDPM)**, progressively transforming Gaussian noise into meaningful latent representations conditioned upon textual input prompts.

The key components of Stable Diffusion include:

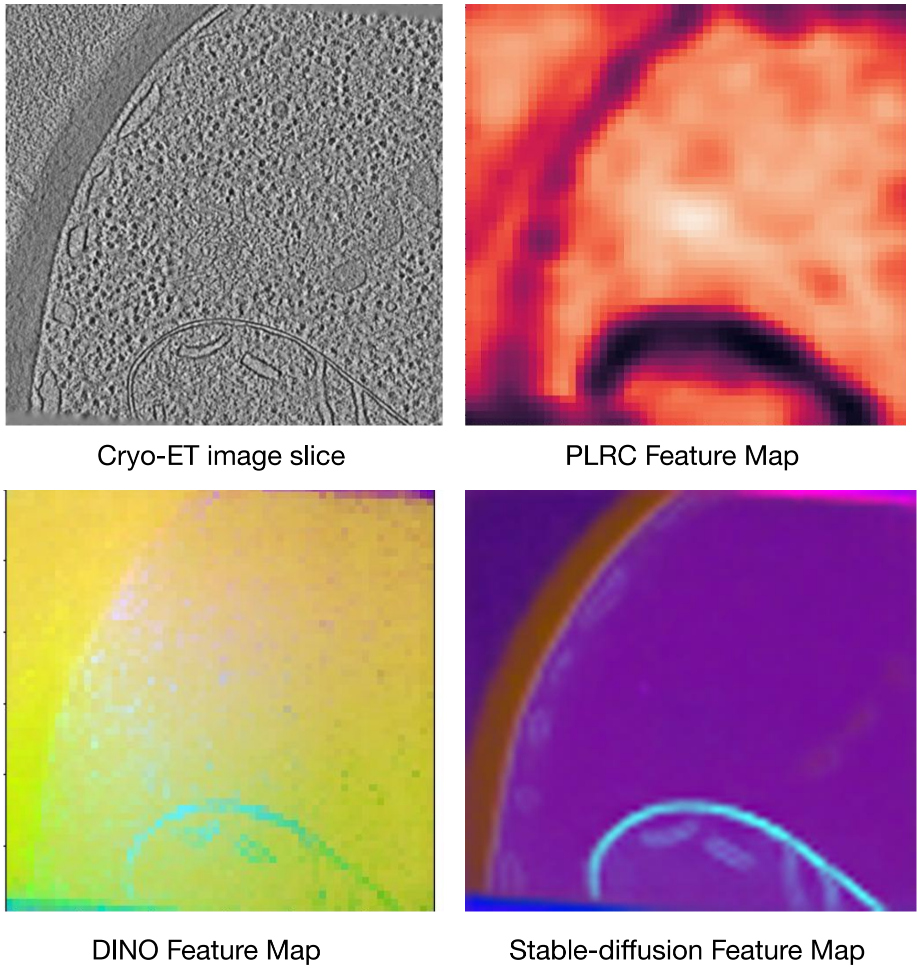


Figure 1: **Feature extraction on a single cryo-ET image slab of a *S. Pombe* Tomogram with multiple VFMs**

- **Variational Autoencoder (VAE):** Encodes high-dimensional images into compact latent representations and subsequently decodes generated latents back into pixel-level images.
- **Text Encoder:** Utilizes a Transformer-based encoder (e.g., CLIP-based) to convert textual inputs into dense embeddings, facilitating conditioning in the diffusion process.
- **UNet Denoiser:** A hierarchical convolutional network combined with Vision Transformer layers, predicting and removing noise in the latent representation space throughout the diffusion steps.
- **Noise Scheduler:** Controls forward (noise addition) and reverse (denoising) processes in latent spaces during training and inference.

In our segmentation pipeline for Cryo-Electron Tomography (Cryo-ET), we leverage the Stable Diffusion model strictly as a frozen feature extractor, without performing any fine-tuning or domain-specific parameter updates. Specifically, we utilize the Vision Transformer backbone within the UNet denoiser to extract dense, high-dimensional features from Cryo-ET images.

Rather than employing Stable Diffusion's generative capacity, we focus on the internal representations obtained from the multi-layer self-attention mechanisms of the ViT backbone. We extract both *query* and *key* embeddings from multiple attention layers, as these embeddings capture rich spatial relationships and structural information essential for accurate unsupervised segmentation.

These embeddings are used to form affinity matrices, characterizing spatial similarity among image patches. Subsequently, spectral clustering is applied to these affinity matrices, allowing eigenvectors to be optimized unsupervisedly to obtain meaningful segmentation masks. By directly using the pre-trained model without modification, our approach efficiently exploits the extensive visual priors embedded in Stable Diffusion, ensuring robust segmentation of complex biological structures in Cryo-ET images.

##### 1.4 Self-attention mechanism

The self-attention mechanism is a key component of vision transformers that enables the model to capture long-range dependencies in an image. Each input image is divided into non-overlapping patches of size $p \times p$, forming a set of tokens $\{x_i\}_{i=1}^{N}$, where $N = \frac{HW}{p^2}$ represents the number of patches, where p = 16. These patches are embedded into a higher-dimensional space and passed through the transformer layers, where self-attention operates as follows:

Each patch embedding $x_i$ is linearly transformed into three vectors:

- **Query** ($Q_i$): Represents the patch's current state in relation to others.
- **Key** ($K_i$): Represents the identifiable characteristics of a patch.
- **Value** ($V_i$): Contains the actual feature representation that will be propagated through layers.

##### 1.5 Affinity Matrix

An *Affinity Matrix* is a fundamental construct in spectral clustering and related unsupervised segmentation techniques, encapsulating pairwise similarities or affinities among elements within a given dataset. In the context of image segmentation, affinity matrices represent the degree of similarity between individual image patches or tokens, quantifying how closely these elements resemble each other in a defined feature space.

Formally, an affinity matrix $A \in \mathbb{R}^{N \times N}$ for a set of $N$ image tokens is defined such that each entry $A(i, j)$ measures the affinity (or similarity) between tokens $i$ and $j$. In our pipeline, affinity matrices are computed based on self-attention mechanisms from Vision Transformer (ViT) layers within the pre-trained Stable Diffusion model.

Specifically, the affinity between tokens $i$ and $j$ for the $l$-th attention layer is computed as:

$$A_l(i, j) = \exp\left(\frac{Q_l(i)K_l(j)^T}{\sqrt{d}}\right),$$

where:

- $Q_l(i)$ and $K_l(j)$ are the *query* and *key* embeddings for tokens $i$ and $j$, respectively, derived from the $l$-th attention layer.
- $d$ is the dimensionality of the embedding vectors.

The exponential function, combined with scaling by $\sqrt{d}$, ensures numerical stability and appropriate normalization of the affinity scores, following the standard practice of scaled dot-product attention mechanisms.

In spectral clustering, affinity matrices serve as a cornerstone for computing eigenvectors that encode meaningful segmentation cues.

##### 1.6 Structural Similarity Index Measure (SSIM)

The Structural Similarity Index Measure (SSIM) is an established perceptual metric that quantitatively assesses the visual similarity between two images by modeling human visual perception more effectively than traditional pixel-based metrics, such as Mean Squared Error (MSE) or Peak Signal-to-Noise Ratio

(PSNR). Rather than solely evaluating pixel intensity differences, SSIM quantifies structural information, capturing similarities in luminance, contrast, and structure between image pairs.

Formally, the SSIM between two image patches, denoted as $x$ and $y$, is defined by:

$$\mathrm{SSIM}(x, y) = \frac{(2\mu_x\mu_y + C_1)(2\sigma_{xy} + C_2)}{(\mu_x^2 + \mu_y^2 + C_1)(\sigma_x^2 + \sigma_y^2 + C_2)},$$

where:

- $\mu_x$ and $\mu_y$ represent the mean intensity values of patches $x$ and $y$, respectively.
- $\sigma_x^2$ and $\sigma_y^2$ are the corresponding variances, capturing local contrast information within each patch.
- $\sigma_{xy}$ denotes the covariance between the patches, capturing the joint variations in their structures.
- $C_1$ and $C_2$ are small stabilization constants to prevent instability when denominators approach zero, typically set as $C_1 = (K_1L)^2$ and $C_2 = (K_2L)^2$, where $L$ is the dynamic range of pixel intensities, and $K_1, K_2 \ll 1$ (commonly $K_1 = 0.01$, $K_2 = 0.03$).

Within our unsupervised segmentation pipeline tailored specifically for Cryo-Electron Tomography (Cryo-ET) images, the SSIM metric is leveraged during the feature refinement phase, particularly in selecting and aggregating structurally coherent eigenvector-based images (eigenimages).

SSIM is strategically employed to evaluate the structural correspondence between candidate eigenimages and the current aggregated feature (base image). Only eigenimages that exhibit a high SSIM with the base image are considered for integration, alongside additional conditions such as improved diversity score and reduced structural noise. This filtering ensures that the merged eigenimages preserve meaningful structures while suppressing noise and irrelevant features.

#### 1.7 Diversity Score

The *Diversity Score* is a heuristic metric developed and utilized within our unsupervised segmentation pipeline to quantitatively assess feature diversity across grayscale eigenvector-derived images. Specifically, the Diversity Score measures spatial variability in pixel intensity distributions, helping to identify eigenimages with rich, informative, and distinct structural content suitable for constructing robust aggregated feature images.

##### Definition and Computation

Given a grayscale image $I \in \mathbb{R}^{H\times W}$, we compute the Diversity Score using the following procedure:

1. **Patch Extraction:** The image $I$ is divided into smaller overlapping or non-overlapping patches of size $p \times p$, with overlap $o$, yielding stride $s = p - o$. Formally, the set of extracted patches $\mathcal{P}$ is defined as:
$$\mathcal{P} = \left\{ I_{ij} \in \mathbb{R}^{p\times p} \;\middle|\; i = 0, s, 2s, \ldots, H - p;\; j = 0, s, 2s, \ldots, W - p \right\}.$$

2. **Local Standard Deviation:** For each extracted patch $I_{ij} \in \mathcal{P}$, we compute the standard deviation of pixel intensities to measure local contrast within the patch:
$$\sigma_{ij} = \sqrt{\frac{1}{p^2} \sum_{(x,y)\in I_{ij}} [I_{ij}(x, y) - \mu_{ij}]^2},$$
where $\mu_{ij}$ is the mean intensity of patch $I_{ij}$.

3. **Aggregation of Local Variances:** The set of local standard deviations across all patches is collected:

$$\boldsymbol{\sigma} = \{\sigma_{ij} \mid I_{ij} \in \mathcal{P}\}.$$

4. **Diversity Score Calculation:** The Diversity Score of image $I$ is defined as the global standard deviation of these local standard deviations, formally given by:

$$\text{DiversityScore}(I) = \sqrt{\frac{1}{|\boldsymbol{\sigma}|} \sum_{\sigma_{ij} \in \boldsymbol{\sigma}} (\sigma_{ij} - \bar{\sigma})^2},$$

where $\bar{\sigma}$ denotes the mean of the local standard deviations in $\boldsymbol{\sigma}$, and $|\boldsymbol{\sigma}|$ is the total number of patches extracted.

Intuitively, a high Diversity Score indicates significant variation in local structural information across the image, which is often indicative of rich feature content and distinct segmentation cues. Conversely, a low Diversity Score implies homogeneity or lack of informative structural details, suggesting that the eigenimage may not substantially contribute to effective segmentation.

Within our Cryo-Electron Tomography (Cryo-ET) segmentation method, the Diversity Score is employed as a critical criterion in selecting and aggregating eigenvector-derived images (eigenimages). Specifically, eigenimages are iteratively evaluated based on their Diversity Scores and structural similarity to construct a robust and informative aggregated feature image. The eigenimage with the highest Diversity Score is selected as the initial 'base' feature, subsequently refined by iteratively incorporating additional eigenimages that simultaneously enhance the Diversity Score and maintain structural coherence. This approach ensures that the final aggregated feature image used for adaptive thresholding captures diverse structural information essential for accurately segmenting biologically relevant subcellular structures.

#### 1.8 Gaussian Adaptive Thresholding

Adaptive thresholding is an effective image processing technique for converting grayscale images into binary images. It dynamically calculates a threshold for each pixel based on the intensities of neighboring pixels, allowing it to adapt effectively to local variations in illumination and texture. The Gaussian adaptive thresholding variant specifically employs a weighted sum of neighborhood pixel values, with the weights defined by a Gaussian distribution.

Formally, Gaussian adaptive thresholding computes the threshold $T(x, y)$ for the pixel located at $(x, y)$ as follows:

$$T(x, y) = \left( \sum_{(u,v) \in \mathcal{N}_b(x,y)} w_{u,v} \cdot I(u, v) \right) - C$$

where:

- $I(u, v)$ is the pixel intensity at location $(u, v)$.
- $\mathcal{N}_b(x, y)$ represents the local neighborhood of size $b \times b$, centered at pixel $(x, y)$. The block size $b$ is typically an odd integer, ensuring a symmetric neighborhood around the center pixel.
- $w_{u,v}$ denotes the Gaussian weight given by:

$$w_{u,v} = \frac{1}{2\pi\sigma^2} e^{-\frac{(u-x)^2 + (v-y)^2}{2\sigma^2}}$$

- $\sigma$ is the standard deviation controlling the spread of the Gaussian kernel.

- $C$ is a scalar constant subtracted from the weighted sum, providing fine control over the thresholding sensitivity and enabling clearer distinctions between foreground and background.

Pixels whose intensity values exceed the computed threshold $T(x, y)$ are classified as foreground (commonly assigned a value of 255), whereas those below the threshold are considered background (commonly set to 0).

This locally adaptive thresholding method significantly enhances detection accuracy and robustness, especially crucial for identifying subtle structural details inherent in Cryo-Electron Tomography (Cryo-ET) imaging.

##### 1.9 UNet architecture

The U-Net architecture implemented here follows the classic encoder–decoder structure, tailored for semantic segmentation tasks. It is composed of two main components: the Encoder, which progressively downsamples the input to capture high-level features, and the Decoder, which upsamples the feature maps to produce a full-resolution segmentation map.

The encoder consists of an initial ConvBlock followed by four DownBlock modules. Each DownBlock applies max pooling to reduce spatial resolution by a factor of two, then passes the result through a ConvBlock consisting of two convolutional layers, batch normalization, LeakyReLU activations, and dropout. The number of feature channels increases at each level, from 16 up to 256, allowing the model to capture increasingly abstract representations. Corresponding dropout rates also increase from 0.05 to 0.5 to provide stronger regularization at deeper layers.

The decoder mirrors this structure, with four UpBlock modules that upsample feature maps using either bilinear interpolation or transposed convolution (in this implementation, transposed convolutions are used). Each UpBlock also performs feature fusion by concatenating the upsampled feature map with the corresponding encoder feature map (skip connection), followed by a ConvBlock. This design enables the decoder to recover spatial detail lost during downsampling. The final segmentation map is produced by a 3×3 convolution that maps the decoder output to the desired number of classes.

##### 1.10 DeepETPicker Training and Inference

DeepETPicker is a deep learning-based method that utilizes a 3D-ResUNet segmentation model with coordinated convolution and multiscale image pyramid inputs to distinguish biological macromolecules from the background in cryo-electron tomograms. The methodology adheres to an established two-stage workflow—training and inference—with identical hyperparameters consistently maintained to ensure methodological rigor.During preprocessing, tomograms undergo coordinate normalization, intensity standardization, and the generation of weak supervision labels—simplified Gaussian-type masks—significantly reducing manual annotation efforts compared to traditional full voxel-level annotations. The training configuration specifies critical parameters: subtomogram size of $72^3$ voxels (chosen to balance memory constraints and spatial context), padding size of 12 voxels for the spatial overlap-based strategy, batch size of 8, learning rate of 0.001, and a segmentation threshold of 0.5. These same coordinates of the same tomogram are also used for validation in this pipeline. During the training phase, subtomograms centered on individual particles are extracted to ensure balanced representation. The 3D-ResUNet model, employing feature maps [24, 48, 72, 108], is trained with these weak labels to maintain accuracy while minimizing annotation costs.In the inference stage, identical spatial parameters are maintained. Tomograms are scanned with subtomogram size N and stride S, following the overlap-tile strategy ($S = N - 2 \times padding_size = 48$ voxels) to eliminate segmentation artifacts at tile edges. A GPU-accelerated pooling-based postprocessing step employing mean-pool non-maximum suppression (NMS) quickly identifies particle centroids from generated segmentation masks.

### 2 Evaluation Metrics

#### 2.1 Membrane Segmentation

To measure the accuracy of membrane segmentation, we used a commonly employed metric to evaluate segmentation methods, known as the Dice Score or Dice coefficient. This score measures the overlap between a predicted segmentation and the ground truth.

Given two sets $A$ and $B$, Dice score is defined as:

$$\text{Dice}(A, B) = \frac{2|A \cap B|}{|A| + |B|}$$

For a 3D voxelized binary segmentation task like ours, $A$ can be regarded as the set of voxels in the ground truth mask and $B$ the set of voxels in the predicted mask.

Then:

$$|A \cap B| = TP, \quad |A| = TP + FN, \quad |B| = TP + FP$$

Substituting into the original Dice score formula:

$$\text{Dice} = \frac{2|A \cap B|}{|A| + |B|} = \frac{2TP}{2TP + FP + FN}$$

We use the final equation to calculate the Dice scores in Table 1 of the main manuscript.

#### 2.2 Macromolecule Localization

To measure the accuracy of macromolecule localization, we used the commonly employed metric, the F1 score. We calculate the F1 score using the list of predicted macromolecule locations and the list of ground truth macromolecule locations. The locations are expressed using the macromolecule's center $(x, y, z)$ coordinate. If any predicted macromolecule location has any ground truth location within 10 voxels of euclidean distance, then it is counted a True Positive (TP). If there are multiple predicted macromolecule location within 10 voxels of euclidean distance to a particular ground truth location, then only 1 of the predicted location is counted as True Positive (TP), while the remaining are counted as False Positive (FP). Any predicted macromolecule location not having any ground truth location within 10 voxels of euclidean distance is counted as False Positive (FP). The ground truth locations that do not have any predicted macromolecule location within 10 voxels of euclidean distance is counted as False Negative (FN). Using the count of TP, FP, and FN, we calculate precision and recall as

$$\text{Precision} = \frac{\text{TP}}{\text{TP} + \text{FP}}$$

$$\text{Recall} = \frac{\text{TP}}{\text{TP} + \text{FN}}$$

.

Finally, the F1 score was calculated as:

$$\text{F1 score} = \frac{2 \times \text{Precision} \times \text{Recall}}{\text{Precision} + \text{Recall}}$$

.
